# The translatome of quiescent *Plasmodium falciparum* gametocytes reveals parasite pyridoxal 5’-phosphate (PLP) biosynthesis is essential for efficient mosquito stage development

**DOI:** 10.64898/2026.03.24.713170

**Authors:** Eduardo Alves, Jack W Houghton, Lindsay B Stewart, Mufuliat T Famodimu, Rosie Bridgwater, Mariana Reis Wunderlich, Naoki Matoba, Annie Tremp, Michaela Bikarova, Jianan Lu, Mojca Kristan, Colin J Sutherland, Edward W Tate, Michael J Delves

**Affiliations:** Department of Infection Biology, London School of Hygiene and Tropical Medicine, Keppel Street, London, WC1E 7HT, UK; LSHTM Malaria Centre, London School of Hygiene and Tropical Medicine, Keppel Street, London, WC1E 7HT, UK; Department of Disease Control, London School of Hygiene and Tropical Medicine, Keppel Street, London, WC1E 7HT, UK; Department of Chemistry, Imperial College London, Molecular Sciences Research Hub, White City Campus, Wood Lane, London W12 OBZ, UK

## Abstract

The ability of *Plasmodium falciparum* gametocytes to remain quiescent within the vertebrate host but poised for rapid onward development in the mosquito is an adaptation essential to maximise the onward spread of malaria. In this dormant state, mature infectious stage V gametocytes are largely unaffected by most antimalarial drugs and our limited understanding of how gametocytes prepare for mosquito transmission has hindered the identification of new molecular targets for transmission-blocking therapeutics. In this study, we move beyond the total proteome of gametocytes and define the translatome of mature stage V gametocytes using L-azidohomoalanine incorporation into nascent proteins, click chemistry purification and proteomic analysis. We identify the proteins and pathways that gametocytes sustain in preparation for transmission during this dormant period and through genetic disruption, we validate this approach by demonstrating the importance of parasite pyridoxal 5’-phosphate biosynthesis for mosquito transmission.

## Introduction

Malaria, a parasitic infection caused by single cell apicomplexan of the genus *Plasmodium*, has shaped both human evolution and civilization for millennia. Despite intense investments and new initiatives aiming to eradicate malaria in the last three decades, it still caused 610,000 deaths in 2024 alone - *P. falciparum* being responsible for more than 90% of mortality cases^1^.

*P. falciparum* cycles between development in the human host and female *Anopheline* mosquitoes. The bite of infected mosquitoes injects sporozoites into human bloodstream where they reach and invade the hepatocytes undergoing an asymptomatic asexual replication ^2^. The infected hepatocytes releases thousands of merozoites that invade the red blood cells (RBC) initiating the intraerythrocytic cycle characterized by ring, trophozoite maturation and schizogony that rupture RBCs resulting on the clinical symptoms of disease^3^. While the majority of merozoites reinvade and restart the intraerythrocytic cycle, some differentiate into sexual forms, the female and male gametocytes. Gametocytes sequester in the bone marrow whilst undergoing 8-10 days of maturation, emerging back into peripheral blood as mature stage V gametocytes ready to infect mosquitoes. Mature stage V gametocytes have reached the pinnacle of their development in the human but have no control over when a mosquito will bite. Therefore, to maximise their probability of successful transmission, they are adapted to live as long as possible whilst maintaining the proteins and pathways essential for their immediate and rapid development into gametes within minutes of entering the mosquito. To achieve this, it is thought that stage V gametocytes enter a quiescent period of limited activity which is reflected in their general insensitivity to most antimalarial drugs designed to target the pathogenic asexual stages^4^. For these reasons, mature stage V gametocytes can escape antimalarial treatment and therefore the “cured” malarial patient can remain part of the infectious reservoir for an extended period^5^.

Given the essential role of gametocytes for mosquito-transmission and their divergence in cell biology from asexual stages, many studies have been performed to identify the protein repertoire expressed by gametocytes ^6–9^. However, all previous studies have focused on the total gametocyte proteome i.e. they describe the collective total of proteins that may have been translated at any point during their lengthy development and accumulated in the mature stage V form. Thus, they do not necessarily provide an accurate snapshot of the biochemical changes and metabolism of the gametocyte just prior to transmission.

In this study, we present an optimised workflow for generating and validating the functionality of pure mature stage V gametocytes. Taking advantage of the high sensitivity of trapped ion mobility spectrometry–time-of-flight (TIMS-TOF) mass spectrometry, we identify an updated and expanded mature stage V gametocyte proteome. Furthermore, for the first time, using metabolic labelling with the azide-bearing methionine surrogate azidohomoalanine (AHA), we identify the translatome of mature stage V gametocytes resolved within a 24 h window. Using gene ontology enrichment analysis, we demonstrate that in preparation for mosquito transmission, mature stage V gametocytes preferentially invest in the translation of distinct proteins and metabolic pathways. Validating this hypothesis, we show that one such upregulated pathway - pyridoxal 5’-phosphate (PLP) biosynthesis is essential for efficient parasite infection and development in the mosquito, thus is an attractive target for transmission-blocking therapeutic interventions.

## Material and Methods

### *In vitro* asexual blood stage parasite culture

*P. falciparum* NF54 strain asexual blood stages were cultured as previously described^10^. Briefly, asexual parasites were kept under 5% parasitaemia (to prevent premature gametocyte commitment) at 4% haematocrit using O, A or AB human erythrocytes (NHS Blood and Transplant Service) under an atmosphere of 1% O_2_/ 3% CO_2_/ 96% N_2_. Culture medium (RPMI-1640, 2.3 g/L sodium bicarbonate, 5.9 g/L HEPES, 50 mg/L hypoxanthine, 1% Albumax II, 5% human AB^+^ serum, 30 mg/ml L-glutamine) was changed daily. Parasitaemia was monitored by blood smear stained with Giemsa. All parasite cultures were confirmed free from *Mycoplasma* contamination via monthly in-house PCR-testing based on a protocol described by Hoppert et al^11^.

### *In vitro* gametocyte culture

Gametocyte cultures were initially seeded from unsynchronized asexual blood stage cultures (4% haematocrit, 2% parasitaemia) and transferred to gametocyte medium (RPMI-1640 with 5% human AB^+^ serum, 0.5% Albumax II, 3.7% 100xHT supplement (ThermoFisher Scientific), 30 mg/ml L-glutamine, 2.78g/L sodium bicarbonate) prewarmed to 37°C at all times (Day 0)^10^. Cultures underwent daily medium changes on a heated surface to maintain culture temperature at 37°C. On Day 5 until Day 9, medium was supplemented with 50 mM of N-acetylglucosamine to remove remaining asexual stage parasites. At Day 14, cultures were predominantly stage V and functional viability confirmed by inducing male gametogenesis (exflagellation) on a small culture sample through temperature decrease and xanthurenic acid stimuli.

### Mature gametocyte purification

Day 15 gametocyte cultures were enriched by density gradient centrifugation using Gentodenz (Gentaur) solution (14.1% w/v Gentodenz, 0.44% w/v NaCl, 0.5 mM Tricine, pH 7.2) in protein LoBind tubes warmed to 37°C (**Figure 1a**). Gametocyte culture was carefully layered on top and centrifuged (800 g, 20 min, 37°C, no brake)^12^. The interlayer enriched for gametocytes was collected and further purified by immobilisation on pre-heated LS magnet-activated cell sorting (MACS) columns (Miltenyi Biotec, UK) attached to a MACS® MultiStand^12^. Columns were washed three times with warm gametocyte medium without human serum and Albumax II to remove non-infected red blood cells (RBC). Purified mature gametocytes were then eluted by gravity flow after detaching the LS columns from MACS® MultiStand holder and quickly pelleted by centrifugation (1000 g, 7 min, 37°C). A Giemsa-stained thin smear was made from the sample to assess gametocyte purity and morphology (**Fig. 1a**). Samples were also tested for the ability to exflagellate to confirm viability. Only purified mature non-activated gametocyte samples demonstrating the ability to exflagellate were used for proteomic experiments. Gametocyte pellets were treated with EDTA-free Halt™ protease inhibitor cocktail (ThermoFisher Scientific) and quickly frozen in dry ice and stored at -80°C.

**Fig. 1.**
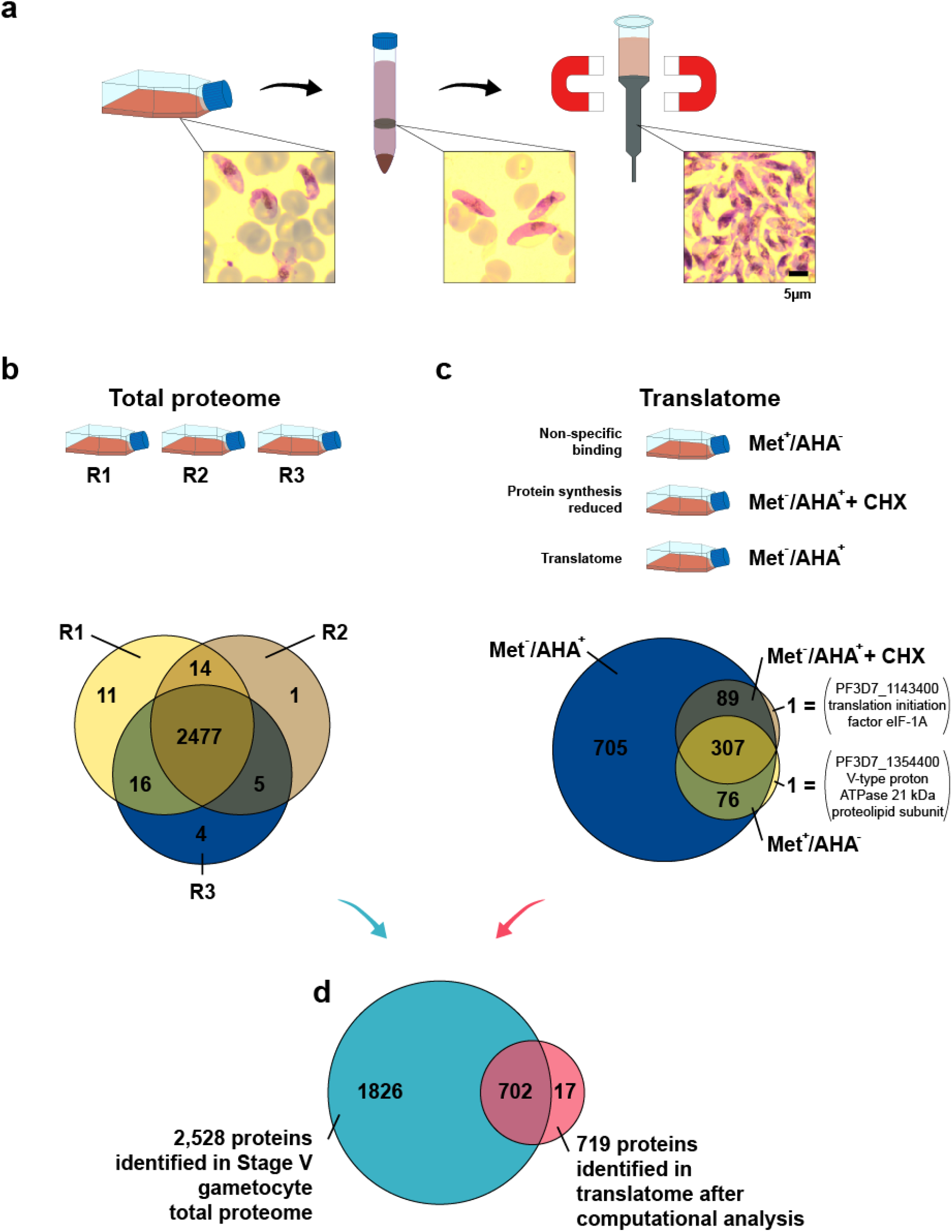
Purification and labelling of *P. falciparum* stage V gametocytes for generating a total proteome and translatome. **a**, Pure stage V gametocytes were obtained using a two-step purification strategy. First gametocytes were enriched by density centrifugation, then collected by magnetic column separation. **b**, A total stage V gametocyte proteome was generated from three independent gametocyte cultures which showed high concordance in identified proteins. **c**, A translatome of stage V gametocytes was generated by incorporation of L-azidohomoalanine (AHA) into nacent proteins over a 24 h period followed by click chemistry-mediated capture and purification. The experiment was performed as a single replicate, but parasites were divided into three groups: the experimental group receiving AHA labelling (Met^-^AHA^+^); a non-specific binding group receiving no AHA (Met^+^ /AHA^-^); and a group receiving AHA labelling but with protein synthesis reduced/inhibited by cycloheximide (Met^-^AHA^+^ + CHX). d, After Significance B analysis, which “rescued” 14 proteins present in the control proteomes but more abundant in the Met^-^ /AHA^+^ proteome, the final stage V gametocyte translatome comprised 27.8% of the proteins of the total proteome.

### Quantification of gametocyte sex ratio

The sex ratio of Gentodenz-purified mature Stage V gametocyte cultures at Day 14 was determined by immunolabelling with an antibody targeting the male-specific marker LDH2^13^. Gametocytes were fixed for 30 min in 4% paraformaldehyde before being washed in PBS and allowed to settle overnight onto poly-D-lysine-coated glass coverslips at 4°C. Fixed cells were then permeabilised with 0.1% triton X-100 for 10 min before being washed in PBS and blocked with PBS + 10% fetal bovine serum (FBS) for 45 min. Anti-LDH2 was diluted 1:10,000 in PBS + 10% FBS and cells stained for 1 h. After washing the cells were finally stained with an anti-rat Alexa488 secondary (1:1,000 in PBS + 10% FBS), washed and mounted with VectaShield™ mountant containing DAPI. The gametocyte sex ratio was quantified by sampling random fields and counting the number of LDH2-positive (male) and LDH2-negative gametocytes (females).

### Quantification of male and female gametogenesis

Gametogenesis was induced in Day 14 gametocytes by reducing culture temperature to 20°C and adding 100 µM xanthurenic acid. A sample of cells were transferred to a Neubauer chamber and 15 min after activation, male gametogenesis (exflagellation) was quantified by brightfield microscopy recording both the number of exflagellation centres and the total number of uninfected erythrocytes. 24 hr later, the remainder of the activated cells were stained live with anti-Pfs25-Cy3 to identify female gametocytes that successfully transformed into female gametes. Cells were transferred to a glass slide and immobilised under a coverslip. Random fields were recorded under brightfield illumination to identify all cells (female gametes and uninfected erythrocytes) and fluorescence to identify Pfs25-positive female gametes^10^. Both male and female gametogenesis were then calculated as a percentage of the total cells in culture including uninfected erythrocytes.

### L-azidohomoalanine (AHA) metabolic labelling of gametocytes

Day 14 gametocyte cultures were pooled (480 ml total volume pooled from 12 × 40 ml cultures) and divided into three groups (160 ml of gametocyte culture each) : 1) A control group that received normal gametocyte medium as previous described above (Met^+^/AHA^-^); 2) A group treated with methionine-free medium (Gibco) supplemented with 5µM of the protein synthesis inhibitor cycloheximide (Sigma Aldrich) for 40 minutes prior to AHA labelling (Met^-^/AHA^+^ + CHX); 3) The experimental group receiving methionine-free medium with 50 µM AHA labelling for 24 h (Met^-^ /AHA^+^).

### Proteome Sample Preparation

The frozen gametocyte pellets were lysed in cold lysis buffer consisting of PBS supplemented with 1% Triton X-100, 0.01% SDS, and EDTA-free Halt™ protease inhibitor cocktail. The lysates were incubated on ice for one hour, then sonicated using an ultrasonic bath (Grant Instruments Cambridge Ltd) at full power for 30 seconds, repeated three times with 30-second intervals. Following sonication, samples were centrifuged at 16,000 × g for 30 minutes at 4°C, and the soluble fraction was collected. Protein concentration was determined using the Pierce BCA Protein Assay Kit. The soluble fractions were either directly processed for whole proteome analysis or used for click chemistry pulldown enrichment.

### Click Chemistry Pulldown

The soluble fraction was incubated at room temperature for one hour on a shaker with a click reaction mix composed of 100 µM PEG4 carboxamide-propargyl biotin (Thermo Fisher Scientific), 1 mM copper sulfate (Scientific Laboratory Supplies), 1 mM of tris(2-carboxyethyl)phosphine hydrochloride (TCEP, Promega UK Ltd), and 100 µM tris((1-benzyl-4-triazolyl)methyl)amine (TBTA, Cambridge Bioscience). The reaction was quenched with 5 mM EDTA. Proteins were precipitated with acetonitrile and resuspended in 50 mM HEPES buffer (pH 8.0) containing 0.2% SDS, followed by incubation with NeutrAvidin agarose beads (ThermoFisher Scientific) for one hour at room temperature on a shaker. The beads were washed four times with the same buffer before proceeding to on-bead digestion.

### Protein digestion

All protein samples, including whole proteome and enriched fractions, were reduced with 10 mM TCEP and alkylated with 40 mM 2-chloroacetamide (CAA), followed by overnight digestion at 37°C using sequencing-grade modified trypsin (Promega UK Ltd).

### LC-MS/MS Acquisition

Peptides (1000 ng) were analysed using an Evosep One (Evosep) system coupled to a timsTOF HT mass spectrometer (Bruker) equipped with a 15 cm × 150 µm, 1.9 µm C18 analytical column (Evosep). Separation was performed using the 30 samples/day (30SPD) workflow with solvent A consisting of 0.1% formic acid (FA) in water, and solvent B consisting of acetonitrile with 0.1% FA. The column was maintained at 40°C. For whole proteome samples, data were acquired in data-independent acquisition mode using parallel accumulation–serial fragmentation (diaPASEF). Acquisition settings included an m/z range of 100 to 1700 and an ion mobility range from 1/K_0_ = 1.30 to 0.85 Vs/cm^2^. Equal ion accumulation and ramp times of 100 milliseconds were used in the dual TIMS analyser. Each acquisition cycle included eight PASEF ramps, covering 21 mass steps with 25 Da isolation windows and non-overlapping ion mobility windows covering 0.85 to 1.27 Vs/cm^2^. Collision energy was dynamically adjusted according to ion mobility, ranging from 59 eV at 1/K_0_ = 1.6 Vs/cm^2^ to 20 eV at 1/K_0_ = 0.6 Vs/cm^2^.

For click chemistry pulldown samples, data were acquired in data-dependent acquisition mode (ddaPASEF), using similar m/z and ion mobility ranges from total proteome samples. Each cycle included four PASEF ramps with a total cycle time of 0.53 seconds, and a 2.75 ms measurement time per precursor. Isolation widths decreased with m/z, from 3 m/z at 800 m/z to 2 m/z at 700 m/z. Active exclusion of previously fragmented ions was applied for 0.4 minutes.

### LC-MS/MS Data Analysis

DIA data from whole proteome samples were processed using DIA-NN (version 1.8.1)^14^ in library-free mode. The analysis used the *Plasmodium falciparum* 3D7 protein database downloaded from UniProt on 15 December 2023, which included 5374 proteins and 246 common contaminants. Deep learning-based prediction of spectra and retention times was enabled. Trypsin specificity was applied, allowing one missed cleavage, with carbamidomethylation of cysteine as a fixed modification and oxidation (M), N-terminal acetylation, and N-terminal excision set as variable modifications, with up to two variable modifications per peptide. Match-between-runs (MBR) was enabled, and quantification was performed using the “Robust LC (high precision)” strategy. Heuristic protein inference was disabled, and both mass and MS1 accuracy were set to 0.

DDA data from the click pulldown samples were analysed using the FragPipe suite (version 5.0.0) with the built-in “LFQ MBR” (Label-Free Quantification with Match-Between-Runs) workflow. Raw .d files were processed with MSFragger (version 3.8)^15^, using strict trypsin digestion with up to two missed cleavages allowed, and a precursor ion tolerance of 20 ppm. Carbamidomethylation of cysteine was set as a fixed modification, while oxidation (M) and N-terminal acetylation were defined as variable modifications. Trimming of N-terminal methionine residues was enabled. The same *P. falciparum* 3D7 database used for DIA-NN was also used here, supplemented with 50% decoy sequences and common contaminants via the inbuilt FragPipe utility. Peptide-spectrum matches were validated using Percolator (version 3.5)^16^, and protein inference and FDR filtering were performed with Philosopher (version 5.0.0)^17^. Label-free quantification was conducted using IonQuant (version 5.0.0)^18^, with match-between-runs enabled and the minimum number of ions for quantification set as 1.

The mass spectrometry proteomics data have been deposited to the ProteomeXchange Consortium via the PRIDE^19^ partner repository with the dataset identifier PXD075878.

### Identification of the stage V gametocyte translatome

Proteins uniquely identified in the AHA-labelled proteome were considered part of the translatome as default. Some identified proteins also were present in the non-specific binding and CHX (cycloheximide)-inhibited control proteomes. To “rescue” proteins from overlapping groups, all spectral data (LFQ) of the translatome, CHX-inhibited control, and non-specific binding control, as well as the LFQ spectral ratios between the translatome and controls, was process using Perseus software^20^. Those proteins that were more abundant in the translatome than other control groups were identified using Significance B analysis after applying the Benjamini-Hochberg procedure for False Discovery Rate (FDR) with a 1% cut off. In total, 14 proteins were rescued using this method.

### Generation of PDX2 KO line

The plasmid pFC-Linker 3 EF1alpha, donated by Prof Ashley Vaughan, containing all elements for CRISPR-Cas9 genome editing, was initially linearized with Esp3I for guide RNA insertion (**Extended Data Fig. 1, Extended Data Table 1**), targeting the first exon of *PfPDX2* (PF3D7_1116200). The repair sequence, containing a codon optimized mNeonGreen with stop codon was designed when integrated to truncate *PfPDX2* to the first 33 bp of the gene. Transfection was achieved using the erythrocyte preloading method. 100 µg of plasmid DNA was dissolved in cytomix (120 mM KCl, 0.15 mM CaCl_2_, 2 mM EGTA, 5 mM MgCl_2_, 10 mM K_2_HPO_4_/KH_2_PO_4_, 25 mM HEPES, pH 7.6) and then used to resuspend 300 µl packed human erythrocytes prewashed in cytomix. The DNA-cytomix-blood was then transferred to an electroporation cuvette and shocked using the single exponential pulse setting (310 V/ 960 µf/ infinite Ω) (Bio-Rad). Electroporated erythrocytes were then immediately transferred into 4.5ml of parasite culture medium and then 0.5 ml of predominantly schizont asexual parasite culture at ∼3 % parasitaemia was added. Cultures were maintained with daily medium changes for 2 days before 5 nM WR99210 was added to the culture medium to select for parasites with episomal expression. Selection pressure was applied for 4 days until no visible asexual parasites were present on a thin Giemsa smear. Then, selection pressure was removed and the parasites allowed to recover (approximately 4 weeks). When parasites recovered, positive selection was applied by treating with 1 µM 5-fluorocytosine to remove parasites still harbouring the episomal plasmid. Resultant parasites were then dilution cloned and successful disruption of *PfPDX2* was confirmed by PCR specific for the genomic integration site (**Extended Data Fig. 1**).

### *In vitro* blood stage asexual growth assay

Sorbitol-synchronized blood stage asexual cultures (as previously described in Lambros et al 1979^21^) at ring stage, from either NF54 wild type or PDX2 KO cultures, were initially split to 0.1% parasitaemia. Each parasite lines were culture in the normal complete asexual medium or complete asexual medium supplement with 0.01g/L (48.5 µM) pyridoxine (vitB6). Samples from all cultures were collected every 24 hours, diluted 5 times with PBS containing the DNA label SYBR safe (ThermoFisher Scientific) 1:5000 and incubated at RT for 5 minutes. Samples were read in a Attune CytPix Flow Cytometer (Invitrogen) using BL1 detector (530/30 nm filter – 488 nm blue laser). RBCs positive for DNA signal was considered infected and a total of 50000 events were counted. Non-infected RBC was used as negative control to delimit negative from positive populations. Parasitaemia growth was monitored until Day 5. Data represent 3 independent experiments with two technical replicates per experiment.

### Mosquito Membrane Feeding

Standard Membrane Feeding Assays were conducted in the LSHTM Human Malaria Transmission Facility (HMTF), as previously described^22^. Day 14 *P. falciparum* NF54 and NF54-PDX2 KO gametocyte cultures showing high levels of stage V gametocytes and high levels of male gametogenesis were diluted to similar gametocytaemias and prepared for blood feeding to mosquitoes. Cultures were pelleted and resuspended with fresh erythrocytes and human serum to ∼40% haematocrit and transferred to water channel-heated membrane feeders. Pots of 70 three-to-five-day-old female *Anopheles coluzzii* (N’gousso strain) mosquitoes were allowed to feed for 30 minutes. Mosquitoes were placed in a climate-controlled incubator at 26.5°C/60% RH/12h ld. Half of the mosquitoes fed with each parasite line were maintained with the standard sugar meal (10% glucose/0.05% P para-aminobenzoic acid (PABA)) *ad libitum*, while the remaining half were given the sugar meal supplemented with 0.01g/L (48.5 µM) pyridoxine (vitB6), which can be taken up by living cells and converted to pyridoxal 5’-phosphate (PLP) via the PLP salvage pathway.

### Oocyst and sporozoite counting and measurements

Midguts from 7-8 days post-fed mosquitoes were dissected and placed on a glass slide with 0.25% mercurochrome stain in PBS. The oocyst intensity (mean number of oocysts per mosquito) and infection prevalence (number of infected mosquitoes) were assessed by phase contrast light microscopy. The same infected oocyst slides were imaged under brightfield illumination using x20 objective lens on a Nikon Ti2 microscope. Oocyst area was measured by drawing the region of interest (ROI) using FIJI (v1.54) software. Salivary gland dissections to record the presence of sporozoites were conducted on infected mosquitoes 14 days post-feed. Glands were dissected and the number of gland lobes collected recorded (each mosquito possesses six lobes but not all are necessarily recovered from each mosquito). All recovered gland lobes for each experimental condition were pooled and homogenised using a glass dounce homogeniser to release intact sporozoites. The volume of homogenate was recorded and then the sporozoite concentration counted in a Neubauer chamber. Using the sporozoite count, volume of homogenate and number of “effective mosquitoes” dissected (number of collected gland lobes/6), the mean number of sporozoites per mosquito was calculated.

### Gene ontology enrichment analysis

The gene IDs for were uploaded to the PlasmoDB (Release 68) website (https://plasmodb.org/plasmo/app) and metabolic pathway enrichment performed using the Kyoto Encyclopaedia of Genes and Genomes (KEGG) as a pathway source, a p value cutoff of p<0.05 and the EC exact match only setting. Metabolic pathways were considered significantly enriched if p<0.05 and was less than the calculated Benjamini-Hochberg false discovery rate (FDR 0.25).

### Statistical Analysis

Unless stated elsewhere, all statistical analyses were performed within GraphPad Prism version 10.

## Results

### The Translatome of *P. falciparum* stage V gametocytes is distinct from the stage V total proteome

To study *P. falciparum* stage V gametocytes in isolation from other parasite stages, we adapted a two-step purification protocol^12^ combining density gradient centrifugation (to remove the majority of uninfected erythrocytes and haemozoin), and then capturing pure stage V gametocytes on magnetic columns (**Fig. 1a**). Stage V gametocytes purified by this method and maintained at 37°C remained non-activated and viable, still retaining a high propensity to form male and female gametes when stimulated by temperature decrease and addition of xanthurenic acid. We applied this purification method to first generate a total proteome of all the proteins present in stage V gametocytes. Strong concordance across three independent replicate gametocyte samples was observed with 98.0 % of identified proteins (2,477) common across replicates and 2,528 proteins identified in total (**Fig. 1b, Extended Data Table 2**). In comparison to previous published proteomes, an additional 426 proteins were identified as expressed in stage V gametocytes (of which 80.3% have previously only been reported as expressed at the transcript level)^23,24^ (**Extended Data Table 3**). To generate a *P. falciparum* stage V gametocyte translatome, gametocytes were treated for 24 h with 50 µM L-azidohomoalanine (AHA), an azide-containing methionine analog readily incorporated into nacent proteins in methionine-free medium (Met^-^/AHA^+^)^25^ (**Fig. 1c**). Gametocytes were purified identically to the total proteome and AHA-incorporated proteins biotinylated through copper-catalyzed azide–alkyne cycloaddition before being captured on strepavidin beads. In parallel, gametocytes were treated identically but incubated either without AHA (Met^+^/AHA^-^) or AHA plus 5 µM cycloheximide (CHX) (Met^-^/AHA^+^ + CHX) to identify proteins non-specifically binding to the streptavidin beads, and to monitor AHA incorporation when protein synthesis is reduced/inhibited respectively (**Fig. 1c**). The Met^-^/AHA^+^ group identified 1,177 proteins, with 705 proteins exclusive to Met^-^/AHA^+^ and only one unique hit in either of the control groups (**Extended Data Table 4**). Interestingly, the unique hit in the Met^-^/AHA^+^ + CHX group was translation initiation factor eIF-1A (PF3D7_1143400). We speculate that this may be a real physiological reponse of the gametocyte to CHX treatment as they attempt to compensate for reduced protein synthesis. To generate the final *P. falciparum* stage V gametocyte translatome, supplementing the 705 proteins exclusive to the Met^-^ /AHA^+^ sample, 14 proteins shared with the control samples were “rescued” using Significance B analysis to determine those common proteins which were significantly enriched in Met^-^/AHA^+^ with a FDR of 1% (**Fig. 1d, Extended Data Fig. 2A**). In total, the translatome comprises 27.8% of the proteins identified in the total gametocyte proteome. In addition, seventeen proteins identified in the translatome were not present in the total protome (**Extended Data Table 5**). Twelve of these have been identified previously in published gametocyte proteomes and sixteen identified in a published gametocyte transcriptome, supporting their Stage V gametocyte identity^23,24^. We hypothesised that the number of methionine residues in a protein could potentially bias the translatome by increasing the likelihood of identification by creating additional biotinylation sites for capture. To test this, we compared the number of methionines per protein with its abdundance in the translatome and found no significant relationship (Pearson’s correlation value = -0.02; p = 0.59) (**Extended Data Fig. 2b**), suggesting that abundance in the translatome is directly related to the relative rate of protein synthesis in Stage V gametocytes.

### The *P. falciparum* Stage V gametocyte total proteome is enriched in proteins associated with protein synthesis and processing, but not cell adhesion and antigenic variation

The twenty most abundant proteins in the Stage V gametocyte total proteome comprised mainly of structural proteins (i.e. beta tubulin, actin 1), glycolytic enzymes (glyceraldehyde 3-phosphate dehydrogenase, enolase, fructose bisphosphate aldolase, phosphoglycerate kinase, lactate dehydrogenase), heat shock proteins, and proteins generally associated with the function of male gametocytes/gametes (lactate dehydrogenase 2^13,26^ and GEST^27^) (**Fig. 2a**).

**Fig. 2.**
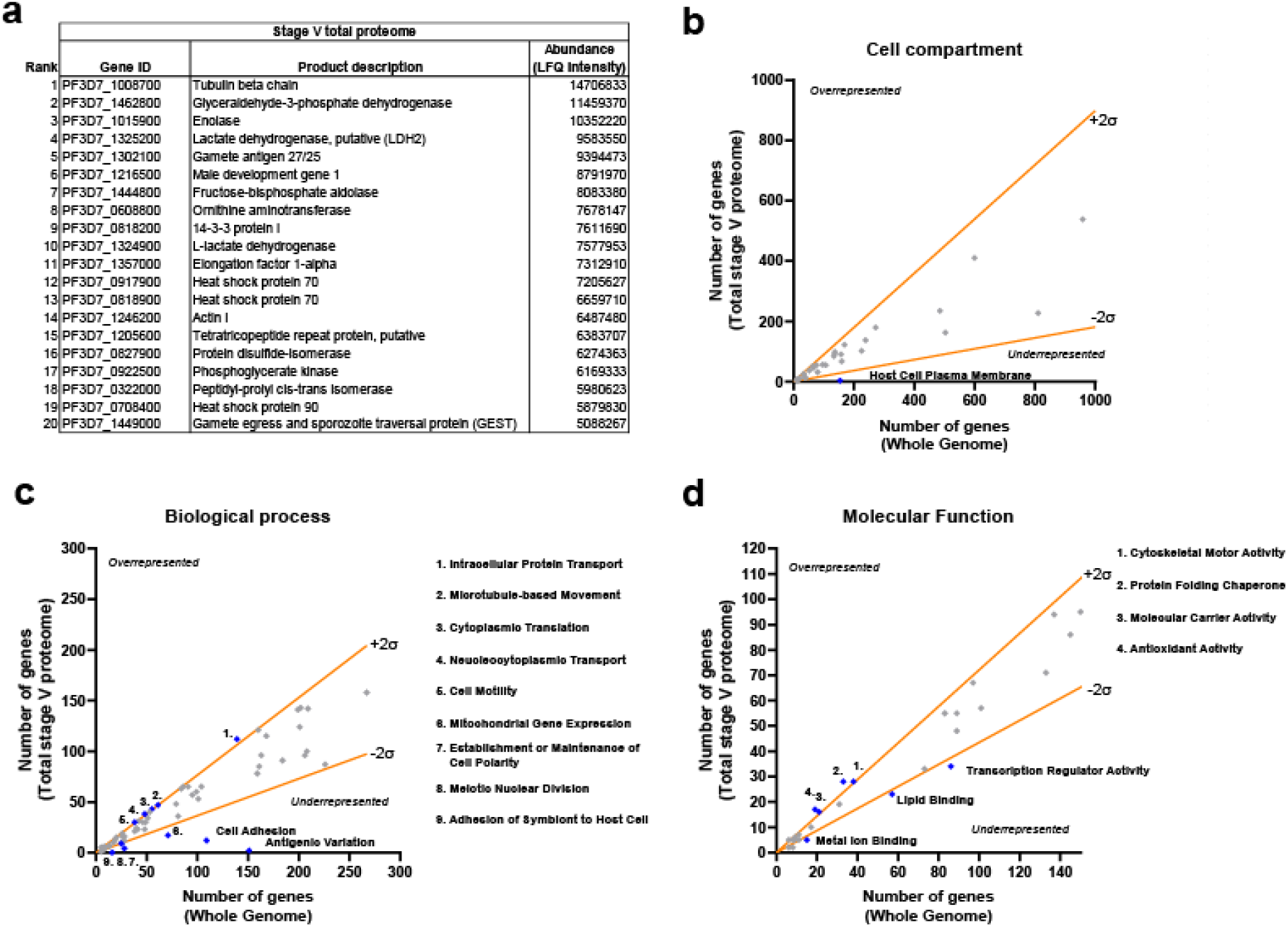
Identity of the most abundant proteins in the total Stage V gametocyte proteome and gene ontology analysis. **a**, The twenty most abundant proteins in the total Stage V gametocyte proteome. Gene ontology (GO) analysis was used to identify GO components over/under-represented in the translatome compared to the total proteome at the **b**, Cell compartment level; **c**, Biological process level; **d**, Molecular function level. Those GO components diverging by ±2 standard deviations from the mean were designated as over/under-represented respectively.

Gametocytes are sexually dimorphic, therefore our total proteome includes proteins present in male or female gametocytes (or both). *P. falciparum* has a frequently reported strong female biased sex ratio (∼0.2-0.35 males per female)^28,29^. Differences in sex ratio between different parasite strains and even strains that have diverged over time in different research groups could substantially affect abundance of differentially-expressed proteins in a total proteome. Therefore sex ratio must be considered when comparing non-sex separated datasets. The NF54 strain used to generate our Stage V gametocyte total proteome was experimentally determined to produce 0.72 males per female (**Extended Data Fig. 3a**). This closer to parity of sex ratio differs from the reported norm, but is not unexpected considering our parasite line has been specifically cultured for years primarily to evaluate male gametogenesis and so would benefit from a higher proportion of male gametocytes. When comparing our proteome to the only sex-separated gametocyte proteome reported^24^, we find that male gametocyte-associated proteins are generally more abundant than expected in our proteome and female gametocyte-associated proteins generally less abundant (**Extended Data Fig. 3b, Extended Data Table 6**), supporting the notion our parasite strain/gametocyte culture methodology generates proportionally more male gametocytes than expected.

Gene ontology (GO) analysis was then used to compare the total stage V proteome to the entire *Plasmodium* genome to give an overview of activity in stage V gametocytes (**Fig. 2b-d, Extended Data Table 7**). A GO term was assigned as overrepresented or underrepresented if the ratio of number of genes (total proteome/whole genome) in that particular GO term diverged more than 2 standard deviations (2SD) from the mean ratio of all the GO terms present. During development, gametocytes are known to export proteins to the erythrocyte membrane that are hypothesised to bind host receptors and assist sequestration in the bone marrow. Upon reaching maturity, these exported proteins are downregulated and removed from the erythrocyte membrane to enable exit from the bone marrow and assist immune evasion in peripheral blood^30^. Supporting this, we find that genes associated with the GO terms “Host Cell Plasma Membrane”, “Cell Adhesion”, “Antigenic Variation”, and “Adhesion of Symbiont to Host Cell” are all underrepresented in mature stage V gametocytes (**Fig. 2b-d, Extended Data Table 7**). Conversely, stage V gametocytes show overrepresentation of GO terms involving protein synthesis, folding and transport within the cell suggesting perhaps investment in preparation of proteins required for current survival and later development. Furthermore, GO terms involved in cell motility are overrepresented, pointing towards the need for male gametocytes to prepare for transmission and rapidly assemble motile male gametes.

### The Translatome of *P. falciparum* Stage V gametocytes is enriched for proteins essential for mosquito transmission

Unlike the total proteome, the twenty most abundant proteins unambiguously assigned to the Met^-^ /AHA^+^ group support a variety of different cellular processes: energy metabolism (i.e. adenylate kinase, α-ketoglutarate dehydrogenase E2 subunit); mitochondrial antioxidant defence (i.e. thioredoxin peroxidase 2, superoxide dismutase); protein translation, trafficking and degradation; and cytoskeletal functions (**Table 1**).

**Table 1.**
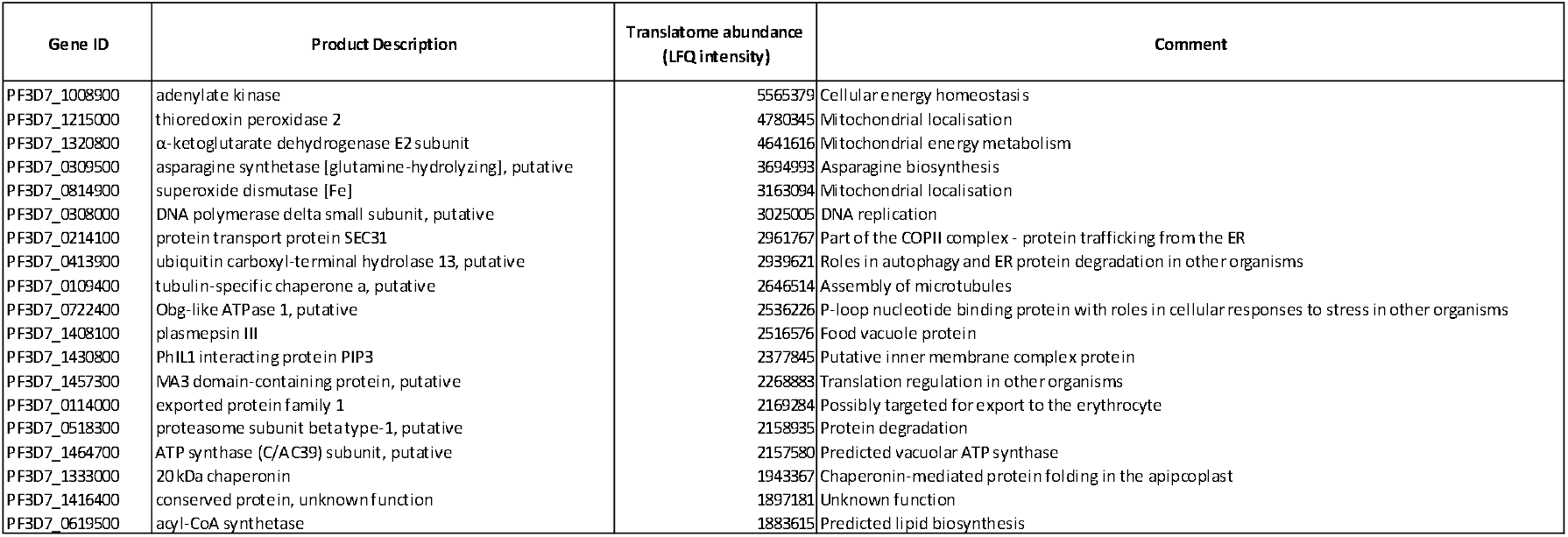
Identity of the most abundant proteins in the Stage V gametocyte translatome. The twenty most abundant proteins identified exclusively in the Met^-^/AHA^+^ condition comprising the Stage V gametocyte translatome.

To further understand which metabolic pathways gametocytes particularly invest in, Kyoto Encyclopedia of Genes and Genomes (KEGG) pathway enrichment analysis was performed for the total stage V proteome and translatome in comparison to the whole *Plasmodium* genome (**Tables 2 and 3**). Strikingly, both datasets highlight B vitamin metabolism as pathways enriched in stage V gametocytes with thiamine (vitamin B1), riboflavin (vitamin B2), nicotinate and nicotinamide (vitamin B3), and pyridoxal 5’-phosphate (vitamin B6/PLP) common to both.

**Table 2.**
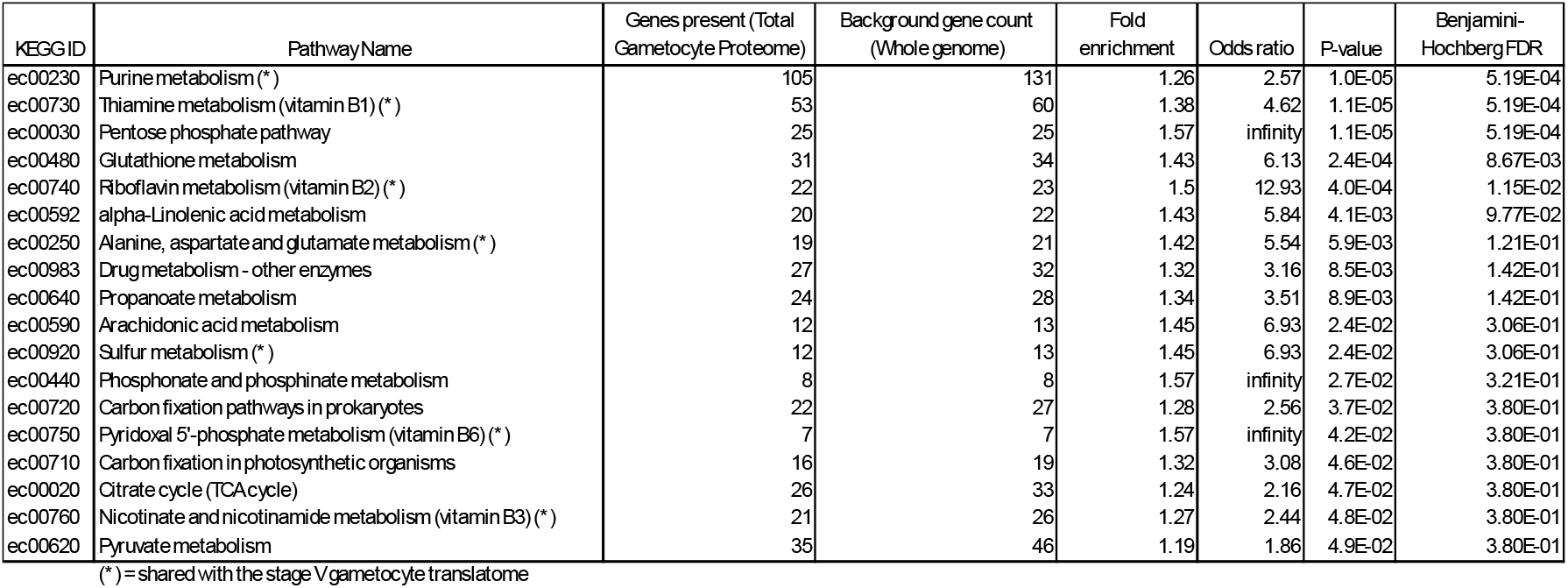
Total proteome metabolic pathway enrichment. The total Stage V gametocyte proteome was analysed by KEGG pathway enrichment analysis, revealing metabolic pathways significantly upregulated in gametocytes compared to the whole genome.

**Table 3.** Stage V gametocyte translatome metabolic pathway enrichment. The Stage V gametocyte translatome was analysed by KEGG pathway enrichment analysis, revealing metabolic pathways significantly upregulated, hence prioritised by the Stage V gametocyte for continual maintenance and preparation for transmission.

| KEGG ID | Pathway Name | Genes present (Translatome) | Background gene count (Whole genome) | Fold enrichment | Odds ratio | P-value | Benjamini-Hochberg FDR |
| --- | --- | --- | --- | --- | --- | --- | --- |
| ec00250 | Alanine, aspartate and glutamate metabolism (*) | 12 | 21 | 2.6 | 4.96 | 4.25E-04 | 4.35E-02 |
| ec00230 | Purine metabolism (*) | 44 | 131 | 1.53 | 2.01 | 6.54E-04 | 4.35E-02 |
| ec00053 | Ascorbate and aldarate metabolism | 11 | 23 | 2.17 | 3.38 | 4.95E-03 | 1.19E-01 |
| ec00740 | Riboflavin metabolism (vitamin B2) (*) | 11 | 23 | 2.17 | 3.38 | 4.95E-03 | 1.19E-01 |
| ec00730 | Thiamine metabolism (vitamin B1) (*) | 22 | 60 | 1.67 | 2.18 | 5.37E-03 | 1.19E-01 |
| ec00565 | Ether lipid metabolism | 22 | 60 | 1.67 | 2.18 | 5.37E-03 | 1.19E-01 |
| ec00981 | Insect hormone biosynthesis | 6 | 10 | 2.73 | 5.45 | 1.00E-02 | 1.91E-01 |
| ec00920 | Sulfur metabolism (*) | 7 | 13 | 2.45 | 4.24 | 1.15E-02 | 1.91E-01 |
| ec00760 | Nicotinate and nicotinamide metabolism (vitamin B3) (*) | 10 | 26 | 1.75 | 2.28 | 4.06E-02 | 5.83E-01 |
| ec00750 | Pyridoxal 5'-phosphate metabolism (vitamin B6) (*) | 4 | 7 | 2.6 | 4.8 | 4.56E-02 | 5.83E-01 |
| ec00470 | D-Amino acid metabolism | 2 | 2 | 4.54 | infinity | 4.83E-02 | 5.83E-01 |

(\*) = shared with the stage V total proteome

Different mass spectroscopy acquisition modes between the total proteome and translatome precluded direct quantitative comparasion of spectal abundance, however relative differences were compared and ranked by dividing their abundance in the total stage V proteome with their abundance in the translatome and applying a Log2 transformation (**Table 4, Extended Data Table 8**). Fifteen proteins showed greater than 2SD increase in Log2(ratio) from the the mean of 1.23 to >4.70 and therefore represent the proteins most synthesised in the translatome compared to their respective abundances in the total proteome. Likely, overrepresentation in the translatome signifies that the protein is labile but must be regularly replaced to maintain viability and onward fertility of the gametocyte. In support of this hypothesis, Rhomboid-like protease 3 (ROM3) and multidrug resistance transporter protein 2 (PfMDR2) were both highly overrepresented in the translatome. The *P. berghei* orthologues of both are reportedly expressed in gametocytes with genetic deletion resulting in lack of sporozoite development later in the mosquito^31,32^. Furthermore, subpellicular microtuble protein 3 (PfSPM3) is reportedly critical for maintaining the falcipform shape of gametocytes indicating an essential role in maintaining gametocyte viability in the human host^33^.

The most overrepresented protein in the translatome was determined to be 60s acidic ribosomal protein P2 (PfP2 - PF3D7_0309600). This ribosomal subunit (along with its counterpart P1) comprises the ribosome stalk region which is thought to play a role in translational regulation in other organisms^34^, but both subunits are rapidly exchanged between the ribosome and exisiting free in the cytoplasm. It would be tempting to speculate that this dynamic exchange could result in a high protein turnover, therefore a need for constant translational replacement. However, PfP1 (PF3D7_1103100), whilst present in the translatome, only ranks 630^th^ in translatome:total proteome abundance suggesting a much lower investment in synthesis of PfP1. Interestingly, PfP2 (but not PfP1) has been shown to have a significant extra-ribosomal role in asexual nuclear division and is exported as a tetramer to the erythrocyte membrane^35^ and we speculate that this may extend also to nuclear division in the mosquito during sporogony.

**Table 4.**
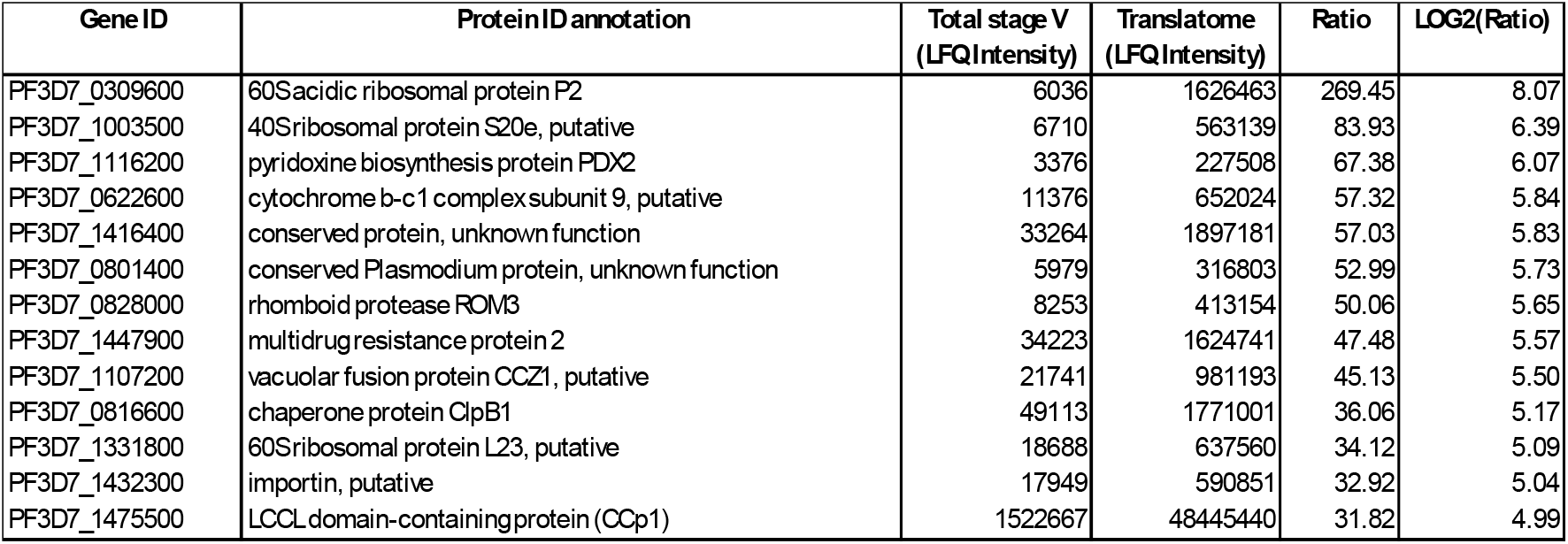
Proteins overrepresented in the translatome. The relative difference in protein abundance was compared between the total Stage V gametocyte proteome and the gametocyte translatome. Thirteen proteins showed greater than 2 standard deviations (2SD) increase in abundance in the translatome.

| Gene ID | Protein ID annotation | Total stage V (LFQ Intensity) | Translatome (LFQ Intensity) | Ratio | LOG2(Ratio) |
| --- | --- | --- | --- | --- | --- |
| PF3D7_0309600 | 60S acidic ribosomal protein P2 | 6036 | 1626463 | 269.45 | 8.07 |
| PF3D7_1003500 | 40S ribosomal protein S20e, putative | 6710 | 563139 | 83.93 | 6.39 |
| PF3D7_1116200 | pyridoxine biosynthesis protein PDX2 | 3376 | 227508 | 67.38 | 6.07 |
| PF3D7_0622600 | cytochrome b-c1 complex subunit 9, putative | 11376 | 652024 | 57.32 | 5.84 |
| PF3D7_1416400 | conserved protein, unknown function | 33264 | 1897181 | 57.03 | 5.83 |
| PF3D7_0801400 | conserved Plasmodium protein, unknown function | 5979 | 316803 | 52.99 | 5.73 |
| PF3D7_0828000 | rhomboid protease ROM3 | 8253 | 413154 | 50.06 | 5.65 |
| PF3D7_1447900 | multidrug resistance protein 2 | 34223 | 1624741 | 47.48 | 5.57 |
| PF3D7_1107200 | vacuolar fusion protein CCZ1, putative | 21741 | 981193 | 45.13 | 5.50 |
| PF3D7_0816600 | chaperone protein ClpB1 | 49113 | 1771001 | 36.06 | 5.17 |
| PF3D7_1331800 | 60S ribosomal protein L23, putative | 18688 | 637560 | 34.12 | 5.09 |
| PF3D7_1432300 | importin, putative | 17949 | 590851 | 32.92 | 5.04 |
| PF3D7_1475500 | LCCL domain-containing protein (CCp1) | 1522667 | 48445440 | 31.82 | 4.99 |

Metabolic pathway enrichment identified vitB6/PLP metabolism as enriched in stage V gametocytes (**Tables 2 and 3**) and pyridoxal biosynthesis protein 2 (PDX2 - PF3D7_1116200) was the third most overrepresented protein in the translatome (**Table 4**). Together with PDX1 (PF3D7_0621200), PDX2 forms the pyridoxal-511-phosphate (PLP) synthase complex. PLP synthase comprises a highly stable dodecamer of PDX1 onto which 12 PDX2 subunits are free to transiently bind^36^. PDX1 and PDX2 are present in the total Stage V gametocyte proteome, but only PDX2 is represented in the translatome. This may reflect differences in protein turnover between subunits with the stable dodecamer PDX1 needing less translational replacement than the dynamic PDX2 subunit. Furthermore, parasites can alternatively salvage PLP via pyridoxal kinase (PLK; PF3D7_0616000)^37^. PLK was also present in the total Stage V gametocyte proteome but absent from the translatome. PLP is an essential cofactor for a wide range of different essential biological pathways^38^. Interestingly, ornithine aminotransferase (PF3D7_0608800), one of the most abundant proteins in the total gametocyte proteome (**Fig. 2a**) requires PLP as a cofactor to function, further highlighting its potential importance to the gametocyte and onward transmission.

### Inhibition of PLP biosynthesis by *pdx2* disruption is detrimental, but not essential for intraerythrocytic development *in vitro* and produces viable gametocytes

To investigate the importance of the unstudied genes and metabolic pathways highlighted in the translatome to parasite mosquito transmission, we focused on PLP biosynthesis as it requires only two genes and cannot be compensated by other pathways apart from the salvage pathway via PLK if exogenous vitB6 is accessible from the host (**Fig. 3a**). Although previous reported attempts to delete *pdx1* and *pdx2* in *P. falciparum* were unsuccessful and chemical inhibition of PXD1 is reportedly lethal to asexual parasites^39^, we noted that piggybac disruption of *pdx2* is reportedly dispensible^40^. We therefore successfully attempted disruption of *pdx2* by CRISPR-Cas9 (PDX2 KO) (**Extended Data Fig. 1**). RPMI medium contains 4.85 µM vitB6 which we hypothesise compensates for *pdx2* loss by via the PLK salvage pathway. Although viable, PXD2 KO showed reduced asexual growth on average 47.6% less than the parental line during sucessive asexual cycles (**Fig. 3b**). Culture medium supplemented with a 10-fold excess of vitB6 (48.5 µM) did not rescue the growth phenotype, suggesting that in the absence of PLP biosynthesis, the parasite PLP salvage pathway is already at maximal capacity and does not fully support growth and development during the asexual cycle.

**Fig. 3.**
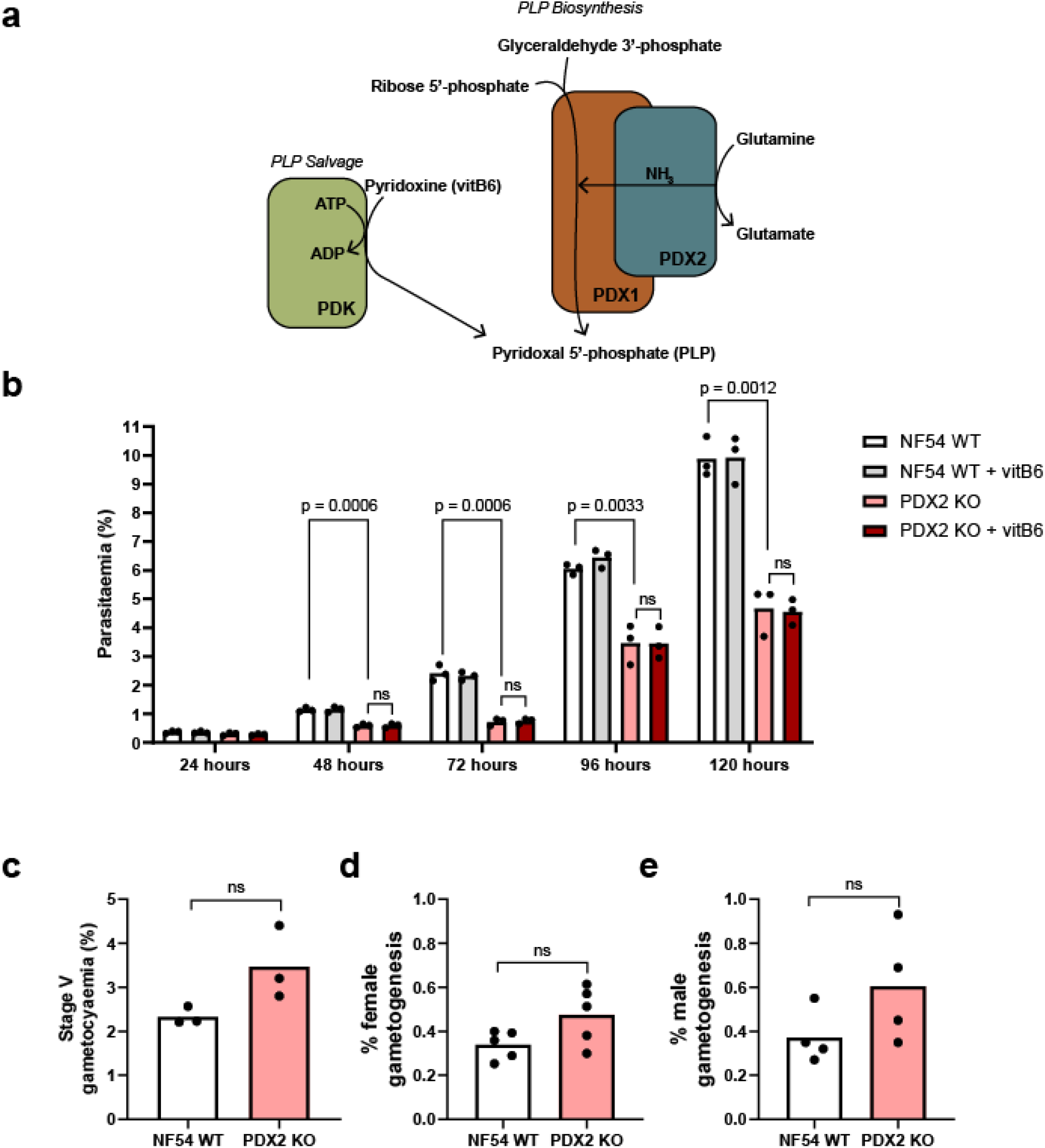
The effect of *pdx2* gene disruption on asexual and sexual parasite development. **a**, Schematic illustrating how *Plasmodium* can obtain pyridoxal 5’-phosphate (PLP) either by biosynthesis via PLP synthase (PDX1 and PDX2) or salvaged from pyridoxine (vitB6) via pyridoxal kinase. **b**, The effect of *pdx2* disruption (PDX2 KO) on asexual parasite growth was compared to the parental wildtype strain (NF54 WT) both with vitB6 concentrations standard for RPMI and ten-fold elevated (+ vitB6). Asexual parasite parasitaemia was quantified every 24 h. Bars show the mean of three independent experiments; black dots show the mean values from two technical replicates of each experiment; p values denote statistical significance using the unpaired Student’s t-test; ns = not significant. The impact of *pdx2* disruption on gametocyte development and functionality was also tested. **c**, Stage V gametocytaemia. Bars show the mean of three independent experiments; black dots show the individual values of each replicate; ns indicates no significant difference as assessed by the unpaired Student’s t-test. **d**, Female gametogenesis. Bars show the mean of five independent experiments; black dots show the individual values of each replicate; ns indicates no significant difference as assessed by the unpaired Student’s t-test. **e**, Male gametogenesis. Bars show the mean of four independent experiments; black dots show the individual values of each replicate; ns indicates no significant difference as assessed by the unpaired Student’s t-test.

Despite a slow asexual growth phenotype, PDX2 KO was able to undergo visibly normal gametocyte development and stage V gametocytaemia was not significantly different from the WT parental line (p = 0.08; unpaired Student’s t-test) (**Fig. 3c**) and showed a similar sex ratio (**Extended Data Fig. 3a**). Furthermore, PDX2 KO gametocytes were demonstratably functional and formed male and female gametes at levels also not significantly different from the WT parental line (p = 0.07 female; p = 0.16 male; unpaired Student’s t-test) (**Fig. 3d+e**). Together, this demonstrates that *in vitro, PDX2* is dispensible for asexual and sexual development even though the gametocyte invests in synthesis of PDX2 protein just prior to mosquito transmission.

### Inhibition of PLP biosynthesis by *pdx2* disruption blocks parasite development in the mosquito and can be rescued by supplementing mosquitoes with exogenous vitB6

Unlike *Plasmodium*, mosquitoes lack a PLP biosynthetic pathway and must salvage PLP primarily through nutritional symbiosis with their gut microbiota^41^ (**Fig. 4a**). Mosquitoes reared under sterile conditions or with antibiotics to clear gut microbiota require vitB6 (and other vitamin) supplementation to develop into viable adults demonstrating its essentiality to the mosquito^42,43^. Therefore, we hypothesised that in the absence of PLP biosynthesis in PDX2 KO, the parasite must compete with the mosquito to salvage available vitB6 and may be at a disadvantage compared to wild type during oocyst development. To study this, we infected *An. coluzzii* mosquitoes with parental WT and PDX2 KO in standard membrane feeding assays (SMFAs) (**Fig. 4b**). Cohorts of mosquitoes were divided in half after feeding and one half were continuously maintained with sugar solution supplemented with 48.5 µM vitB6 and the other half without. Across four independent biological replicates, PDX2 KO showed a significant 53.8% decrease in median oocyst intensity compared to the parental WT line (6 oocysts per mosquito vs 13 oocysts per mosquito respectively; p = 0.007 by Mann-Whitney test with Bonferroni correction for multiple comparisons) which was rescued by addition of exogenous vitB6 to the sugar solution of PDX2 KO-infected mosquitoes (p = 0.0002). A significant reduction in infection prevalence (percentage of infected mosquitoes) was not observed across any of the experimental conditions (**Fig. 4c**). Given the observed parental WT oocyst intensity under our experimental conditions and an observed 53.8% reduction of intensity with PDX2 KO, this is to be expected with the reported saturating relationship between oocyst intensity, prevalence and efficacy of transmission reducing interventions^44^. However although present, in the absence of exogenous vitB6, at Day 7 after infection PDX2 KO oocysts were 40.1% smaller than those of the WT parental line, showing a median area of 287 and 424 μm^2^ respectively (p < 0.0001 by Mann-Whitney test using Bonferroni correction for multiple comparisons). This was also rescued by vitB6 supplementation (p < 0.0001). Interestingly, vitB6 supplementation of the parental WT line reduced oocyst area by 21.8% (p < 0.0001). We hypothesise that this reduction may reflect increased competition for resources or overcrowding of oocysts in WT parental-infected mosquitoes which could limit their capacity to grow given that the vitB6-supplemented mosquitoes had a generally higher (but not significantly higher) oocyst intensity. Together, our data demonstrates that a deficiency in parasite PLP biosynthesis not only affects the infection efficiency of the parasite to the mosquito, but also impairs its continuing development within the mosquito which is directly linked to available PLP.

**Fig. 4.**
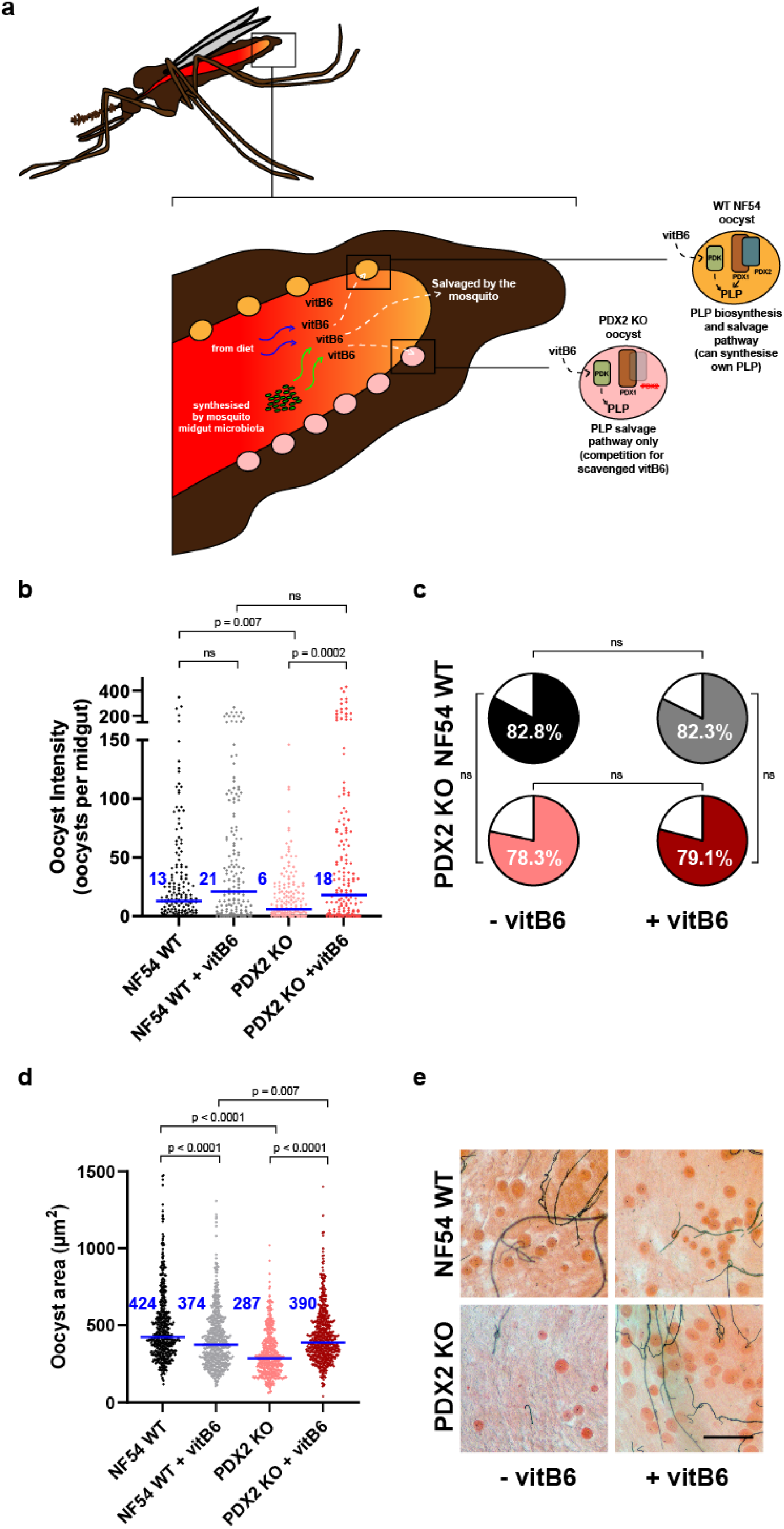
The effect of *pdx2* disruption on parasite transmission to the mosquito in a standard membrane feeding assay (SMFA). **a**, Diagram illustrating exogenous sources of pyridoxal 5’-phosphate (PLP) available to the parasite in the mosquito gut either derived from dietary ingestion of pyridoxine (vitB6) by the mosquito or synthesised by the mosquito midgut microflora. WT parental parasites (WT NF54) may also synthesise PLP via PLP synthase. PDX2 KO parasites must rely on PLP derived from the salvage pathway. **b**, Oocyst intensity of parental WT (WT NF54) infected mosquitoes compared to PDX2 KO either maintained on glucose solution or glucose containing 48.5 µM pyridoxine (+ vitB6) for seven days after infection. Data shown is the combined outcome of four independent biological replicates (n = 147-157 mosquitoes dissected per condition; blue lines and text indicate median oocyst intensity values; p values indicate significance using Mann-Whitney test using Bonferroni correction for multiple comparisons). **c**, Infection prevalence of parental WT (WT NF54) infected mosquitoes compared to PDX2 KO either maintained on glucose solution (- vitB6) or glucose containing 48.5 µM pyridoxine (+ vitB6) for seven days after infection. Data shown is the combined outcome of four independent biological replicates (n = 147-157 mosquitoes dissected per condition; conditions compared using the two-sided Fishers exact test for statistical significance. **d**, The area of individual oocysts from each condition was measured and compared. Data shown is the combined outcome of two independent biological replicates (n = 522-555 oocysts; blue lines and text indicate median oocyst area; p values indicated significance calculated by Mann-Whitney test using Bonferroni correction for multiple comparisons). e, Mercurochome-stained mosquito midguts showing representative oocyst size. Images shown are taken from the same biological replicate (bar = 100 µm).

To explore this further, we asessed how the reduction in oocyst intensity of PDX2 KO oocysts combined with their reduced size impacted subsequent salivary gland sporozoite load (**Fig. 5, Extended Data Table 9**). In two independent biological replicates, we observed that disruption of *PDX2* resulted in a 60.6-90.4% reduction in salivary gland sporozoite burden (from 10,087 sporozoites per mosquito to 968 sporozoites per mosquito in replicate 1 and 2052 to 809 sporozoites per mosquito in replicate 2) which was also rescued by continuous post-feed supplementation of mosquitoes with 48.5 µM vitB6 (11,761 sporozoites per mosquito in replicate 1 and 3099 in replicate 2). Sporulating midgut oocysts were readily observed in the parental WT-infected mosquitoes and also both WT and PDX2 KO infected mosquitoes supplemented with vitB6. However, few sporulating oocysts were observed in PDX2 KO infected mosquitoes without supplementation and those observed were substantially smaller than WT. When combined with the oocyst phenotype, loss of PLP biosynthesis continues to impede parasite development in the mosquito, likely ultimately resulting in a substantial reduction in parasite infectivity to subsequent human hosts.

**Fig. 5.**
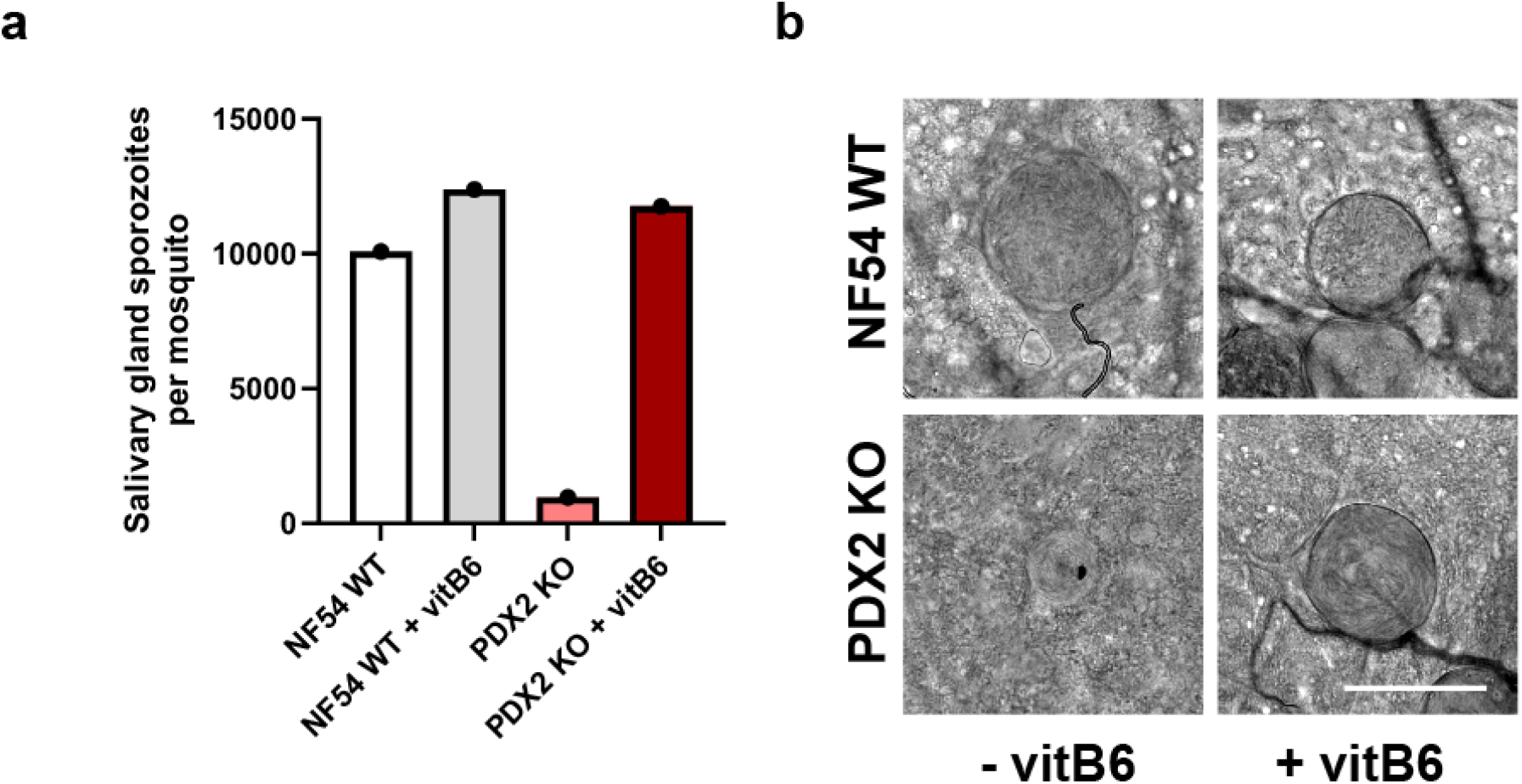
The effect of *pdx2* disruption on salivary gland sporozoite burden and sporulation. **a**, Mean number of salivary gland sporozoites per mosquito of parental WT (WT NF54) infected mosquitoes compared to PDX2 KO either maintained on glucose solution or glucose containing 48.5 µM pyridoxine (+ vitB6) for 14 days. Data shown is one biological replicate with an additional replicate presented in Extended Data Table 8 (n = 26-28 mosquitoes dissected per condition). b, Differential interference contrast microscopy of midgut sporulating oocysts taken from the same mosquitoes used for sporozoite quantification. Bar = 50 µm.

## Discussion

*P. falciparum* is peculiar amongst human-infective malaria species in that gametocyte development is prolonged and once maturity is reached, they become metabolically less-active. Therefore, it is difficult to disentangle whether the proteome of the mature gametocyte is mainly derived of proteins translated earlier in development, and what proportion of the proteome is translated on an ongoing basis once it reaches maturity and infectivity. In order to progress in its life-cycle and propagate, *Plasmodium* mature stage V gametocytes must be ready for uptake and onward development in the mosquito – a sudden, singular event with no second chances. Therefore, to maximise their probability of transmission success, gametocytes must both maintain themselves in the human host as long as possible, and carry a full complement of proteins required for successful transformation and establishment of infection in the mosquito. By specifically studying the translatome of gametocytes upon reaching maturity, our data captures the proteins and pathways that maintain gametocyte viablity and infectivity.

Our metabolic labelling of nascent proteins translated within a 24 h period gives a snapshot of activity within the gametocyte and shows that mature stage V gametocytes still are substantially translationally active. Likely limitations in methodology and purification methods result in not all nascent proteins being captured, nevertheless, our dataset puts a lower limit on the stage V gametocyte translatome of 27.8% of the total stage V gametocyte proteome. Interestingly, through comparative and GO enrichment analyses, we find that particular genes and specific biochemical pathways are prioritised for translation. Notable amongst our datasets, we find that gametocytes prioritise translation of proteins involved in multiple B vitamin biosynthesis and salvage pathways. Like humans, mosquitoes lack biosynthetic pathways for B vitamins and must acquire them from their diet or symbiotically through their gut microflora^41,45^. Therefore, the ability of of *Plasmodium* to synthesize its own B vitamins *de novo* is presumably an essential adaptation to ensure parasite survival in the mosquito where availability of microbial or dietary-derived B vitamins is uncertain or intermittent. Vitamin B5/pantothenate is already a validated antimalarial drug target with MMV693183 (a pantothenate antimetabolite) showing transmission-blocking activity against female gametocytes^46^. Taking advantage of these validated links between B vitamins and mosquito transmission, our study instead focused on vitB6/PLP. We demonstrate that loss of the parasite PLP biosynthesis pathway through genetic disruption of *Pfpdx2* reduces both asexual development and development of the parasite in the mosquito. The former is not rescued by exogenous vitB6, and the latter completely rescued by exogenous vitB6. This does not appear to be due relative differences in ability to salvage PLP via PLP kinase as transcriptomic studies have found that PLP kinase transcript levels are broadly similar between asexuals and oocysts and it is only upgegulated once the parasite reaches sporozoite stage^47^. Therefore, we hypothesise that asexual parasites have high demand for PLP in a short window of time (completing an entire asexual cycle including invasion, growth and replication within 48 h) and that PLP salvage may saturate before these needs are completely met. Contrastingly in the mosquito, after ookinete penetration of the midgut wall, oocyst development to sporogony in the mosquito is relatively slow, occuring over 12-18 days. We speculate that whilst also essential in the mosquito, PLP salvage can accumulate PLP over a longer period of time and so in the presence of excess exogenous vitB6, the parasite can better tolerate loss of PLP biosynthesis.

In the absence of excess exogenous PLP, PDX2 KO parasite infectivity and development in the mosquito was cumulatively reduced, with a reduction in oocyst numbers, a reduction in oocyst area and subsequent reduction in salivary gland sporozoite numbers. This suggests that the parasite is scavenging some PLP from the mosquito – either directly or indirectly from its gut microflora, but that it is not adequate to restore full infectivity. At first glance, this may indicate that antimalarials designed to target PLP synthase may be less effective than predicted and therefore not optimal for transmission-blocking therapies. However, this may acutally open new opportunities for both malaria and mosquito control. Small molecules that target both PLP synthase in *Plasmodium* and in the mosquito microbiota could act to block parasite transmission and development by preventing the parasite from both synthesising its own PLP and accessing PLP from the gut microbiota. Previous studies have linked B vitamin depletion to reduced mosquito fecundity and survival^48–50^. Therefore, such an approach also targeting the gut microbiota would limit PLP availability to the mosquito itself, and so could form the basis of a new mosquito control intervention either impregnated onto bednets, or added to attractive targeted sugar baits. In this manner, by targeting parasite transmission, parasite development in the mosquito (either to eliminate or slow/reduce sporogony) and reducing the lifespan and fecundity of the mosquito, interventions targeting PLP could provide a multifaceted approach to breaking the cycle of malaria transmission for which the barrier to evolution of resistance may be too steep.

By demonstrating the importance of PDX2 for parasite transmission, we validate our translatome as containing proteins with essential functions for the gametocyte and onward parasite development in the mosquito. Therefore, mining this dataset may be an efficient starting point for studying parasite adaptations for mosquito transmission and ultimately the identification of candidates for target-based screening campaigns for transmission-blocking antimalarial drug discovery.

## Funding

This project was supported by a UKRI MRC Career Development Award (MR/V010034/1) and Medicine for Malaria Venture Grant (RD-21-1003) awarded to MJD, a Wellcome Biomedical Resources grant (221363/Z/20/Z) awarded to CJS.

## Author Contributions

The project was conceived by MJD, EWT and EA. EA prepared all parasite material for proteomics with guidance and support from JWH, JL and EWT. Proteomic analysis was performed by EA, JWH and EWT. Transgenic parasite generation was performed by EA, with culture assistance from MTF, LS and RB. Mosquito transmission experiments were performed by the LSHTM Human Malaria Transmission Facility (CJS, MK, LS, MRW, MB, AT, NM, MJD)

## Competing Interests

The authors declare no competing interests.

---

**Extended Data Figure 1.**
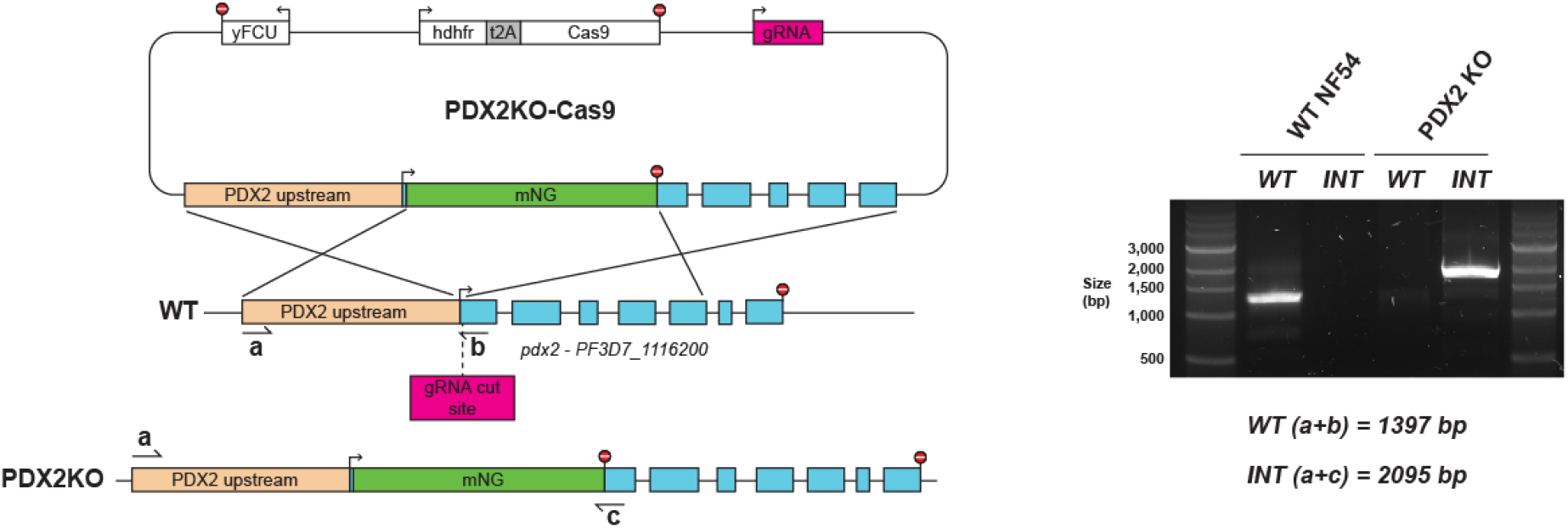
Development of the PDX2 KO parasite line. *pdx2* was disrupted by CRISPR/Cas9 to insert mNeonGreen containing a stop codon into the first exon of the gene. Diagnostic PCR using primers specific to the wildtype (WT) locus and modified locus (INT) were used to confirm successful modification and clonality of PDX2 KO.

**Extended Data Figure 2.**
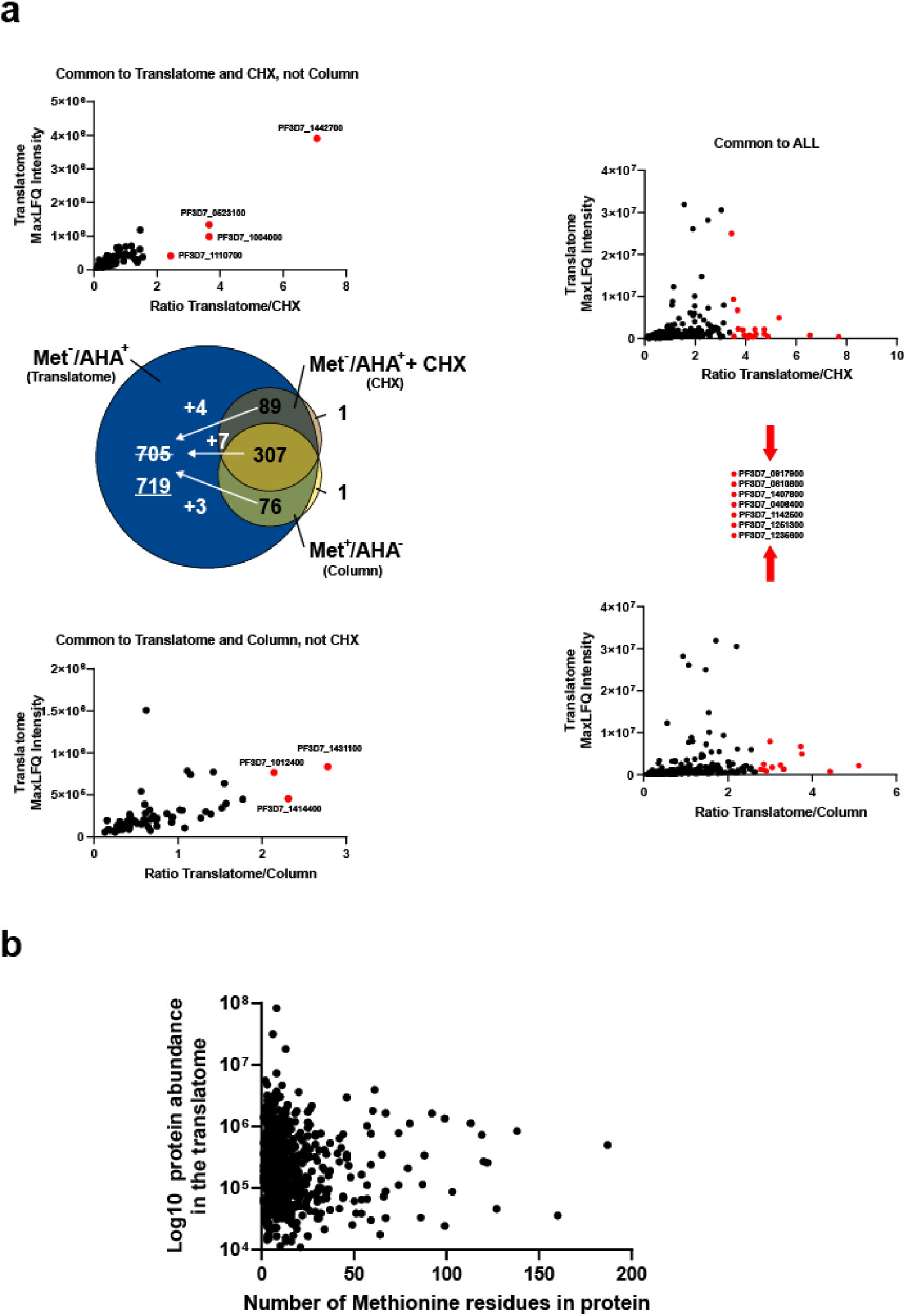
Refining the *P. falciparum* mature stage V gametocyte translatome through significance B analysis and studying the relationship between presence in the translatome and protein methionine content. **a**, Significance B analysis with 1% false discovery rate cut off was used to “rescue” proteins common in both the Met^-^/AHA^+^ dataset and the control datasets, but significantly enriched in Met^-^/AHA^+^. In total fourteen proteins were rescued. **b**, In order to confirm capture and identification in the translatome dataset was not related to the abundance of methionine residues in the protein, the relationship between protein abundance and methionine residues was studied. No significant relationship was demonstrated (Pearson’s correlation value = -0.02; p = 0.59).

**Extended Data Figure 3.**
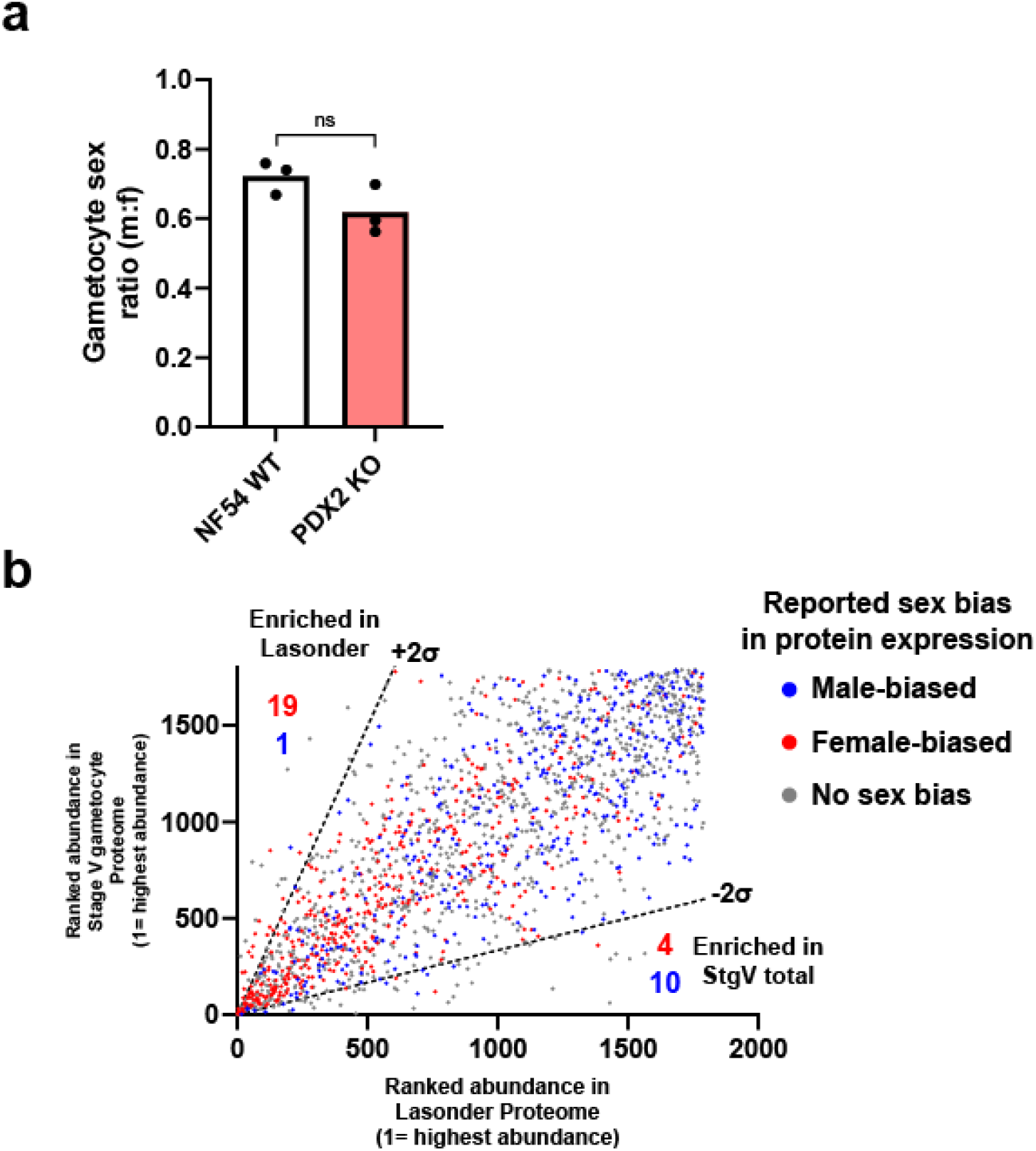
Exploring the effect of gametocyte sex ratio on the total stage V gametocyte proteome. **a**, The gametocyte sex ratio of the parental NF54 strain used in this study was closer to parity than has been more frequently reported in the literature that usually reports ∼0.2-0.3 m:f sex ratio. After genetic modification to disrupt *pdx2*, gametocyte sex ratio of the transgenic parasite was not significantly different from WT (p = 0.11 as calculated by the unpaired Student’s t-test; n = 3 independent experiments).

## Notes

### Competing Interest Statement

The authors have declared no competing interest.

