## Extended Data Table 1 for "The translatome of quiescent *Plasmodium falciparum* gametocytes reveals parasite pyridoxal 5’-phosphate (PLP) biosynthesis is essential for efficient mosquito stage development": Extended Data Table 1.docx

| Sequence  5’-3’ | Description |
| --- | --- |
| catattaagtatataatattatagggg  tattatcactacaaggtagagctagaa  atagcaagt | Forward primer for gRNA integration by Gibson cloning. |
| acttgctatttctagctctaccttgtagt  gataatacccctataatattatatactt  aatatg | Reverse primer for gRNA integration by Gibson cloning. |
| tgcggttcaaaatcacgttgtta  tccgtcgactttatacaattcatc | Forward primer to amplify mNeonGreen and complement PDX2 repair sequence. |
| aactataggggtattatcactaatg  gcaagtttgccagcaac | Reverse primer to amplify mNeonGreen and complement the PDX2 5’ homology region for Gibson cloning. Also used to confirm genome integration (Extended Data Figure 1A - Primer C). |
| acaacgtgattttgaaccgcatataa  atcattttatcaaattacaaatacc | Forward primer to amplify the PDX2 repair sequence, introduce a point mutation to eliminate the PAM site for the gRNA. |
| tagctatgtcgactatccgcgtacttc  atctgataatatttctcttatataagg | Reverse primer to amplify the PDX2 repair sequence and complement the CRISPR plasmid for Gibson cloning. |
| agtgataatacccctatagttatttctgac | Reverse primer to amplify the 5’ homology region. |
| atagcaaaagaaaagaaagcggta  ttatcataatatccagaagcac | Forward primer to amplify the 5’ homology region and complement the CRISPR plasmid for Gibson cloning. Also used to confirm assess either WT genotype or PDX2KO integration (Extended Data Figure 1A - Primer A) |
| gttcaaaatcaccttgtagtgataatac | Primer targeting only the native WT *pdx2* sequence. Used to confirm WT genotype (Extended Data Figure 1A - Primer B) |
| CACTTGTGCAGGTTGTATTCTC | Forward primer spanning the exon 3-4 boundary of the *pdx2* transcript. Used to assess loss of *pdx2* in the PDX2KO parasite (Extended Data Figure 1B - Primer X). |
| CAATTATTCTGTTCAACGGCTGC | Forward primer spanning the exon 6-7 boundary of the *pdx2* transcript. Used to assess loss of *pdx2* in the PDX2KO parasite (Extended Data Figure 1B - Primer Y). |
